# Corpus callosal microstructure predicts bimanual motor performance in chronic stroke survivors: A preliminary cross-sectional study

**DOI:** 10.1101/2021.05.14.443663

**Authors:** Rini Varghese, Brianna Chang, Bokkyu Kim, Sook-Lei Liew, Nicolas Schweighofer, Carolee J. Winstein

## Abstract

Microstructural changes in the corpus callosum are associated with more severe motor impairment in the paretic hand, poor recovery, and general disability. Considering its role in bimanual coordination, we suspected that these microstructural changes across the callosum may also be reflected in the performance of ecologically valid routine bimanual tasks. Thus, the purpose of this study was to determine if callosal microstructure predicts bimanual motor performance in chronic stroke survivors by examining the regions of the corpus callosum connecting both the sensorimotor and non-sensorimotor cortices. We examined the relationship between the fractional anisotropy across the CC and movement times for two self-initiated and self-paced bimanual tasks in 41 chronic stroke survivors. Using publicly available control datasets (n = 52), matched closely for acquisition parameters, we also explored the effect of stroke and age on callosal microstructure. There were two main findings: First, callosal microstructure was significantly associated with bimanual performance in chronic stroke survivors. Notably, a significant relationship was observed not only with the primary sensorimotor regions, but also regions of the premotor/supplementary motor and prefrontal regions. Second, chronic stroke survivors presented with significantly lower mean FA, compared to neurologically intact adults. We conclude that in mild-to-moderate chronic stroke survivors with relatively localized lesions to the motor areas, callosal microstructure can be expected to change in not only the primary sensorimotor region, but also more anteriorly in the secondary motor regions and the genu and is associated with performance on cooperative bimanual tasks.

**Significance:** A goal of rehabilitation after stroke is to promote the return to pre-stroke levels of upper limb function and use, predominantly characterized by coordinated bimanual activities. In this study, we find that in the chronic phase of stroke, microstructural disorganization within the corpus callosum predicts motor performance on real-world bimanual tasks and lends important insight into the indirect, remote effects of stroke.

**Highlights:**

- We provide initial evidence that corpus callosal microstructure predicts performance on two self-initiated and self-paced bimanual tasks.
- Associations were strongest for fibers connecting the primary sensorimotor cortices followed by the pre- and supplementary motor, and prefrontal cortices.
- Significant reductions in fractional anisotropy were observed in stroke survivors for all regions of the corpus callosum.

## Introduction

Focal ischemic injury to the central nervous system can result in changes remote from the site of injury (*diaschisis*, ^1^). One such case is *transcallosal* diaschisis in which the ischemic event in the lesioned cortex triggers structural and functional alterations in its contralateral homolog through the corpus callosum (CC). In recent years, noninvasive imaging of the CC microstructure, e.g. using diffusion tensor imaging-derived metrics like the fractional anisotropy (FA) index, has proven useful in predicting motor recovery after stroke ^2–5^ as well as response to physical and occupational therapy.^6^

These studies have revealed important insights regarding the status and evolution of microstructural changes in the CC following a stroke. For example, recently, Pinter and colleagues showed that callosal microstructure can be expected to change as early as 72 hours post-stroke, not only in the primary sensorimotor regions, but also the non-primary sensorimotor regions.^7^ These changes, particularly disorganization of fibers in the anterior callosum or genu, indexed by lower FA, were found to be associated with general disability and predicted motor recovery at 3 months.^7^ Persistence of lower FA in the callosum in the chronic phase has also been correlated with paretic hand motor impairment and function.^8^

However, these studies were limited by small sample sizes and an almost exclusive focus on traditional clinical measures of unilateral (paretic) motor impairment or disability, which may only weakly correspond to changes in CC microstructure, particularly its non-sensorimotor regions. Conversely, based on the long-established evidence for the role of CC in interlimb coordination ^9–15^, lower FA in the non*-*sensorimotor CC regions might be better reflected in the performance of *bimanual* tasks that preferentially engage bi-hemispheric circuits.

To address these limitations, the primary purpose of this study was to determine if CC microstructure predicts bimanual motor performance in chronic stroke survivors by examining fibers connecting both the sensorimotor and non-sensorimotor regions. Two candidate regions of the callosal genu that were of special interest for the control of bimanual skills were the prefrontal region (CC1), involved in higher-order planning and response selection ^16,17^, and, the premotor and supplementary motor regions (CC2), involved in temporal sequencing. ^18–20^ We hypothesized that lower FA in not only the primary sensorimotor but also CC1 and CC2, would correspond with poor performance on the bimanual task.

In a secondary exploratory analysis, we assessed the presence of FA reductions across the CC by comparing our own data in chronic stroke survivors (n = 41) to a publicly available dataset of neurologically intact, age-similar (n = 24), and younger (n = 28) control adults. This control dataset allowed us to more cleanly isolate the effects of stroke from the more general effects of age, establishing a baseline from which the stroke effects can be compared. Based on Pinter et al and Hayward et al., we hypothesized that compared to age-similar controls, FA would be significantly reduced, beyond the general reductions from aging, in all regions of the CC, including the non-sensorimotor regions.

## Methods

### Participants

Diffusion tensor imaging data for 41 chronic stroke survivors were available from a Phase 2B randomized controlled trial (Dose Optimization for Stroke Evaluation, ClinicalTrials.gov ID: NCT01749358) (Winstein et al., 2019). Only baseline data from the DOSE study were included in this analysis. These data were collected between 2012 and 2015 on the Health Sciences Campus of the University of Southern California (USC).

For our exploratory analysis examining differences in the microstructural status of the CC in chronic stroke survivors versus healthy controls, we used publicly available diffusion datasets acquired in 24 age-similar older adults and 28 younger adults, matched closely for acquisition parameters, including sequence, diffusion gradient strength and number of directions (OpenNeuro.org ID: ds001242). These data were collected between 2016 and 2018 on the University Park Campus of USC.

All individuals gave informed consent to participate in the two studies in accordance with the 1964 Declaration of Helsinki and the guidelines of the Institutional Review Boards of the respective campuses of USC where the data were collected.

### Diffusion Imaging

#### Acquisition

The diffusion MRI scans in stroke survivors was acquired on a GE Signa Excite 3T scanner using a single shot spin echo EPI pulse sequence with the following parameters: TR = 10,000 ms, TE = 88 ms, FoV = 256 mm, 75 axial slices of thickness = 2.0 mm, gradient strength of 1000s/ mm ^2^ in 64 diffusion gradient directions. This generated a 2 × 2 × 2 voxel size and a matrix size of 128 × 128. A high-resolution structural T1-weighted image was acquired prior to diffusion imaging using the gradient-echo (SPGR) sequence with the following parameters: TR = 24 ms, TE = 3.5 ms, flip angle = 20 □, FoV = 240 mm, and slice thickness = 1.2 mm with no gaps, generating a matrix size of 197 × 233 × 189. Total time was approximately 20 minutes.

The younger and age-similar older control datasets were acquired on a Siemens 3T Trio Total imaging matrix (Tim) system. A comparison of all scanner acquisition parameters is provided in the Supplementary Material (I).

#### Preprocessing

Data pre-processing and analysis followed a standard pipeline using the FMRI Software Library, FSL (Figure 1). Voxel-wise statistical analysis of the FA data was carried out using TBSS (Tract-Based Spatial Statistics, Smith et al., 2006), part of FSL (Smith et al., 2004). Diffusion images were first preprocessed, including correction for eddy currents and motion-related distortion, followed by brain extraction (BET2, Jenkinson et al., 2005). Next, FA images were created by fitting a tensor model to the preprocessed diffusion data using FDT (Smith, 2002). All subjects’ FA data were then aligned into a common space (FMRIB58_FA) using the nonlinear registration tool FNIRT (Andersson et al., 2007a, 2007b), which uses a b-spline representation of the registration warp field (Rueckert et al., 1999). The FMRIB58_FA is a white matter template generated from an average of 58 high-resolution, well-aligned FA images from healthy adults and is in the same coordinate space as the 1mm MNI 152 template. Visual quality checks (QC) were performed at the end of every step to ensure accurate registration and tensor fitting. Lastly, the mean FA image was created and thinned to render an FA skeleton which represents the centers of all tracts common to the group. Each subject’s aligned FA data was then projected onto this skeleton and a voxel-wise threshold FA of 0.2 is applied to remove any edge effects.

**Figure 1.**
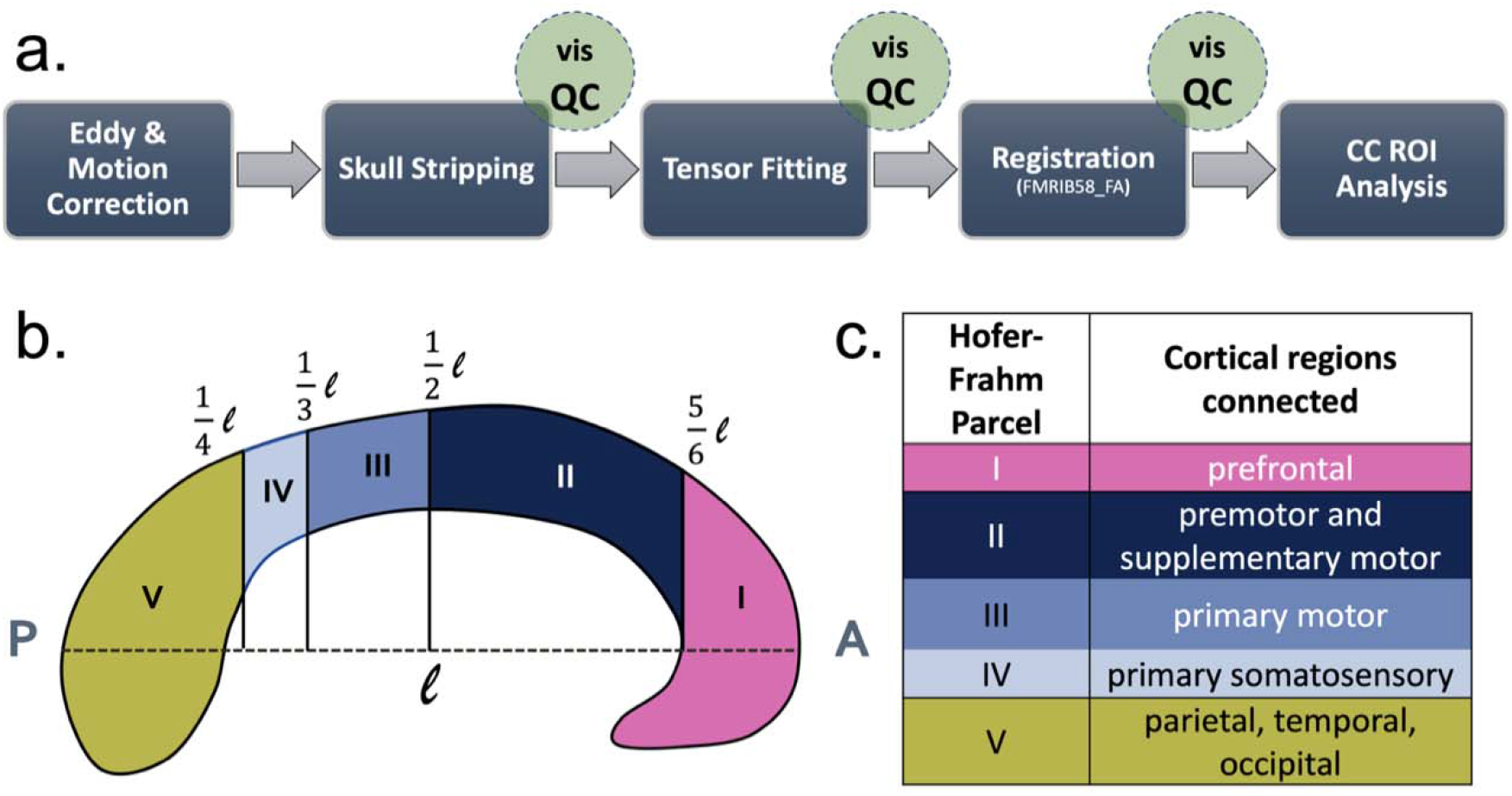
a) Diffusion pipeline including preprocessing, co-registration and identifying region of interest (i.e., corpus callosum). (b) Parcellation (from posterior to anterior) of the corpus callosum using the geometric scheme proposed by Hofer & Frahm (2006). (c) Table showing cortical regions connected by the fibers running through each of the five CC segments.

### Assessment of Callosal Microstructure

The corpus callosum (CC) was defined as the primary region of interest. To assess microstructural status of the CC, we analyzed diffusion images in the standard space, and used the JHU white matter atlas to mask the CC (JHU ICBM-DTI-81 White-Matter Labels). The CC was then segmented geometrically on the midsagittal plane into five subregions according to the Hofer-Frahm parcellation scheme (Hofer & Frahm, 2006). Each of these segments correspond to fibers connecting homotopic regions of the prefrontal (I), premotor and supplementary motor (II), primary motor (III), primary somatosensory (IV), and parietal, temporo-occipital (V) cortices, which in the standard MNI space, consists of 8851, 6871, 5803, 2619, 12729 voxels respectively. Mask templates are available in the first author’s OSF repository: <u>osf.io /7j9xe</u>

Microstructural status was quantified as the fractional anisotropy (FA) index. The FA index is a composite measure reflecting the 3-dimensional directional characteristics of diffusion in each voxel, serving as a proxy for fiber orientation (Hagmann et al., 2006; Pierpaoli & Basser, 1996; Soares et al., 2013). It is computed as a normalized fraction of the eigenvalues derived directly from voxel-wise fitted tensors, and ranges from 0 (isotropic diffusion, spherical in shape) to 1 (anisotropic diffusion, ellipsoidal in shape). The FA composite measure works particularly well for directionally homogenous, well-aligned fibers such as those of the CC, especially after thresholding for edge effects. In a random subset of stroke survivors (n = 20), we validated the FA index generated in the standard-space CC mask with those in the native FA maps and found no difference in mean FA between the two spaces. Results from these comparisons and bootstrap analyses are provided in Supplementary Materials (II).

In stroke survivors only, we also computed tissue volume as an index of CC macrostructure. To do this, individual CC masks were drawn in the native space of each participant’s structural T1 image using ITK-SNAP (v. 3.8). We normalized CC volumes to express them as a percentage of total white matter volume. To compute total white matter volume, we performed tissue segmentation using FSL’s FAST routine with visual quality checking to ensure that all viable white matter tissue, sparing the lesion, was identified in the segmentation procedure.

### Lesion Reconstruction

Stroke lesions were manually drawn on structural T1 images by trained personnel using MRIcron, an open-source tool for segmentation and visualization (Rorden & Brett, 2000). A detailed procedure has been described previously and all T1-weighted images and binarized lesion masks are available as part of the ATLAS stroke lesion database, vR1.1 (Liew et al., 2017). Lesion volume was calculated using FSL’s *fslstats* function. A lesion overlap image among stroke survivors was generated using the *fslmaths -add* function and visualized in FSLeyes (Jenkinson et al., 2012).

### Bimanual Motor Performance

In conjunction with diffusion imaging, behavioral data for 33 of the 41 right-handed stroke survivors were available for analysis (see Supplementary Material II for a full description of all 41 participants). The behavioral paradigm has been described in detail previously (Varghese et al., 2020). Briefly, participants were covertly observed as they performed the letter-envelope task of the Actual Amount of Use Test. The letter-envelope task consisted of two components: folding the letter then inserting the letter into the envelope.

Data were captured on video and analyzed offline to quantify whether participants chose a unimanual or bimanual strategy and the time taken to complete the task at self-selected speed, i.e., movement time. Start times were defined as the frame when initial contact was made with the letter or envelope, and end times were defined as the frame when the goal was accomplished, i.e., when the last fold was completed, or letter was fully inserted into the envelope. Movement time (MT) was defined as the time elapsed between the start and end time points and was determined for each component of the composite task.

Given that the strategy was self-selected, the speed was self-paced, and the testing itself was conducted unbeknownst to the participants, performance on this task was largely unconstrained, serving as a proxy for interlimb coordination in those who chose a bimanual strategy, as if it were in the real world, even if qualitatively variable between individuals.

### Statistical Analysis

All analyses were conducted using the R statistical computing package (version 3.5.1). All continuous variables, age, chronicity, Upper Extremity Fugl-Meyer scores (UEFM), and movement time, were assessed for normality. A one-way ANOVA was used to compare age among the three groups (i.e., younger controls, older controls, and stroke survivors) followed by pairwise comparisons using Tukey’s HSD. Kruskal-Wallis test was used to compare the proportion of females and males among the three groups. Significance was set at p = 0.05 and adjusted for multiple comparisons when necessary.

Distributions for chronicity and movement time were positively skewed and so they were log-transformed. Assumptions for generalized linear models, including linearity, equality of variance, independence and normality of errors were met and model diagnostics, including leverage and multicollinearity of independent variables, were tested when appropriate.

#### Relationship between callosal microstructure and bimanual performance in chronic stroke survivors

First, in chronic stroke survivors only, to determine if mean callosal FA can be used to predict MT, we used linear mixed effects regression of the form below:

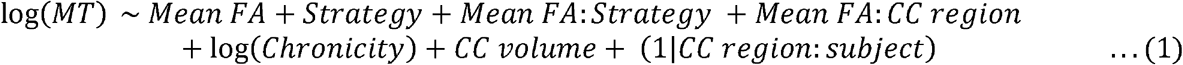

To confirm that the above hypothesized relationship between mean CC FA and MT is in fact due to the coordinative elements of bimanual performance rather than an epiphenomenon emerging from weakness of one limb, we needed to rule out such a relationship between CC FA and unimanual performance. To do this, we tested for a moderating effect of strategy on the relationship between FA and MT (mean FA x strategy, where strategy was coded as bimanual, 0 or unimanual, 1). Marginal slopes for each strategy were then tested for significant difference from 0 using Tukey’s HSD. A significant relationship in those who chose a unimanual strategy would suggest that performance on this task is not compromised due to transcallosal diaschisis alone but at least in part due to the motor capacity of the affected hand.

To test our a-priori hypothesis that lower FA in not only the primary sensorimotor but also non-sensorimotor regions (CC1, prefrontal & CC2, premotor and supplementary motor) would correspond with poor performance, we included a term to test the moderating effect of CC region on the relationship between mean FA and MT. CC region was a categorical variable with 5 levels to code for the segments of the CC, with CC3 (motor) set as the reference level.

Because a single value for MT per subject was repeated over five CC regions, there was no reason to suspect MT to change as a function of CC region. There may be, for instance, an additive shift in the random variance associated with subject and regional differences in intercepts, e.g., mean FA value for CC5 could be higher than CC3 in subj# 1 but lower in subj# 5. Thus, to estimate variance from this additive shift, we modeled the random effects as an interaction between subject and CC region. Once again, marginal slopes for each CC region were then tested for significant difference from 0 using Tukey’s HSD.

To arrive at the final model (eq. 1) we used a combined—forward then backwards— stepwise approach, in which we tested for the confounding effects of age, sex, chronicity, side of lesion, UEFM score, and normalized total CC volume by adding them to the base model that consisted only of mean FA, strategy, and CC region. Then, from the combined model, we removed predictors that were not significant (p < 0.05). Based on this selection process, only log-transformed chronicity and normalized total CC volume were included in the above final model. Notably, by including normalized total CC volume, we were able to consider the likely loss in CC tissue volume.

#### Comparing callosal microstructure between chronic stroke survivors and neurologically intact adults

Second, to explore the effect of stroke on mean callosal FA, we used linear mixed effects regression of the following form:

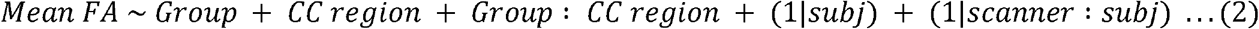

Pairwise comparisons of estimated marginal means for each CC region were conducted using Tukey’s HSD. Our hypothesis was that mean FA would be lower in stroke survivors compared to age-similar adults. We suspected that while group effects would be largest for CC3 (motor) directly adjacent regions (e.g., CC2, premotor) would also show significant reductions in FA. Given that our data were obtained from two different scanners, random effects were estimated as random intercepts for both subject- and scanner-related variances.

Here again, we arrived at the final model (eq. 2) through the same process described above, testing for the confounding effects of age and sex; neither met a cut-off *p* = 0.1, so were removed from the above final model. Note that age was in fact partially embedded within the grouping factor itself. However, because testing for age-related effects on FA was not the primary purpose of this study, we only preserved age as a categorical variable. A supplementary analysis of the relationship between age and FA is provided for the interested reader (Supplementary Material IV).

## Results

Table 1 provides demographic information. Chronic stroke survivors consisted of 22 individuals with left hemisphere stroke and 19 with right hemisphere stroke. There was no significant difference in age (*p* = 0.196), sex (*p* = 0.529), chronicity (*p* = 0.409), or UEFM (*p* = 0.633) between the two stroke groups. Lesion volume was slightly larger in those with right hemisphere strokes but not significant (Δ mean = 1945.4 cc, *p* = 0.054).

**Table 1.** Subject characteristics.

|  | CONTROLS |  | STROKE<br>(N=41) |
| --- | --- | --- | --- |
|  | Younger<br>(N=28) | Older<br>(N=24) |  |
| **Age (years)** |  |  |  |
| Mean (SD) | 24.4 (5.07) | 67.0 (5.55) | 59.1 (13.1) |
| **Sex** |  |  |  |
| Female | 9 (32.1%) | 9 (37.5%) | 11 (26.8%) |
| Male | 19 (67.9%) | 15 (62.5%) | 30 (73.2%) |
| **Chronicity (years)** |  |  |  |
| Median [Min, Max] |  |  | 1.90 [0.474, 14.4] |
| **UE Fugl-Meyer (/66)** |  |  |  |
| Median [Min, Max] |  |  | 43.0 [19.0, 58.0] |
| **Lesion Volume (cc)** |  |  |  |
| Median [Min, Max] |  |  | 6.05 [0.0160, 121] |

On average across all stroke survivors, the lesion constituted < 0.05% (∼11 voxels) of the total CC volume, whereas voxels of the CC constituted < 0.2% (∼2 voxels) of the total lesion volume, confirming a very minor degree of direct injury to the CC. Figure 2 shows lesion overlap among 41 chronic stroke survivors. Individual descriptions of lesion locations are provided in the Supplementary Material (III).

**Figure 2.**
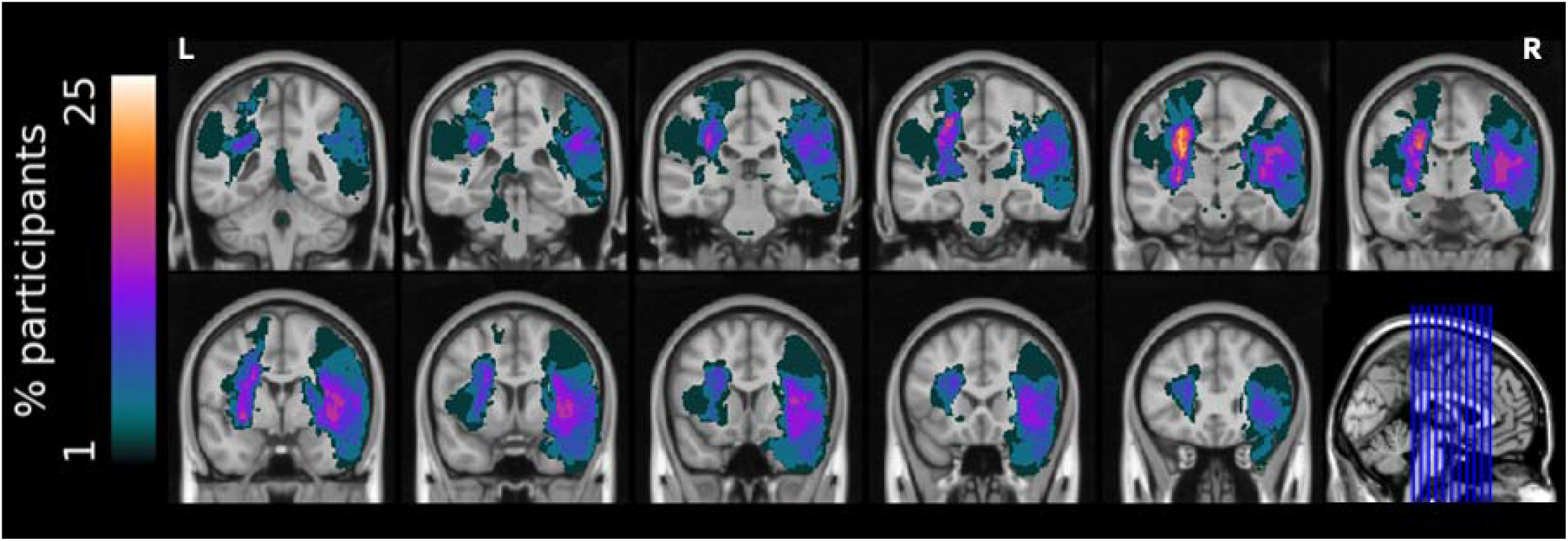
Overlap of lesions for 41 chronic stroke survivors. Note that images were not flipped and represent the actual side of unilateral stroke.

### Result 1: Lower callosal FA is associated with slower bimanual but not unimanual performance in chronic stroke survivors

We found that after accounting for chronicity and total normalized CC volume, mean FA significantly predicted movement time in chronic stroke survivors who selected a bimanual strategy; that is, lower (more isotropic) FA was associated with slower performance. Table 2 provides model estimates from mixed-effects regression. To interpret values in Table 2, please note again that the reference level for strategy was ‘bimanual’ whereas that for CC region was ‘CC3’ (motor).

**Table 2.** Robust mixed-effects regression coefficients from model (1) to estimate relationship between movement time and mean fractional anisotropy (FA), moderated by strategy as well as the five segmented regions of the CC. Note again that CC3 (motor) was the reference level (thus the factor ‘Mean FA’ is the slope for CC3 and estimates for other levels are added to this estimate to derive individual slopes presented in the post-hoc marginal trends).

| *Predictors* | **log(mt)** |  |  |
| --- | --- | --- | --- |
|  | *Estimates* | *CI* | *p* |
| Intercept | 5.05 | 4.05 – 6.05 | **<0.001** |
| Mean FA | -3.60 | -5.15 – -2.06 | **<0.001** |
| Strategy | -1.87 | -3.96 – 0.21 | 0.078 |
| log(Chronicity) | 0.16 | 0.08 – 0.23 | **<0.001** |
| Total Normalized CC Volume | -0.06 | -0.19 – 0.06 | 0.337 |
| Mean FA x Strategy | 2.82 | 0.01 – 5.62 | **0.049** |
| Mean FA x CC1 | 0.27 | -0.04 – 0.59 | 0.092 |
| Mean FA x CC2 | 0.13 | -0.18 – 0.43 | 0.418 |
| Mean FA x CC4 | 0.33 | -0.00 – 0.66 | 0.053 |
| Mean FA x CC5 | 0.61 | 0.20 – 1.01 | **0.003** |
| **Random Effects** |  |  |  |
| $\sigma^2$ | 0.28 | | |
| $\tau_{00}$ CC_region:subjID | 0.03 | | |
| ICC | 0.10 |  |  |
| N<sub>CC_region</sub> | 5 |  |  |
| N<sub>subjID</sub> | 33 |  |  |
| Observations | 330 |  |  |
| Marginal R<sup>2</sup> / Conditional R<sup>2</sup> | 0.144 / 0.227 |  |  |

Post-hoc tests of marginal trends revealed that slope was significantly different from 0 for those who chose a bimanual strategy (*b* = -3.34 ± 0.72, p < 0.001), but not for those who chose a unimanual strategy (*b* = -0.52 ± 1.55, p = 0.74). As expected, marginal slope was largest for CC3 (motor, *b* = -2.19 ± 1.03, *p* = 0.035), followed closely by CC2 (premotor, *b* = -2.07 ± 1.07, *p* = 0.041) and CC1 (prefrontal, *b* = -1.92 ± 0.97, *p* = 0.05). The slope for CC3 did not significantly differ from CC1 and CC2 as observed in the interaction terms, Mean FA x CC1 and Mean FA x CC2. This suggests that consistent with our hypothesis, FA of premotor and prefrontal CC were both similarly predictive of bimanual MT as the motor CC. The slopes, however, were less steep for CC4 (sensory, *b* = -1.87 ± 0.96, *p* = 0.053) and CC5 (parietal, temporo-occipital, *b* = -1.59 ± 0.9, *p* = 0.079) as observed in the interaction terms, Mean FA x CC4 and Mean FA x CC5. However, post-hoc comparisons of all slopes revealed that the slope of only CC5 was significantly smaller than CC3 (*t* = -2.93, *p* = 0.031).

### Result 2: Compared to neurologically intact adults, chronic stroke survivors exhibit lower FA in all regions of the CC, except the splenium

There was a significant interaction between group and CC region (*F* (8, 360) = 21.01, *p* < 0.001). Compared to neurologically intact older adults, mean FA was lower for all CC regions, except the splenium (parietal, temporo-occipital region). Greatest decrements were seen for the primary motor region, CC3 (ΔFA = 0.052, *t* = 4.84, *p* < 0.001), but was closely followed by the premotor and supplementary motor, CC2 (ΔFA = 0.050, *t* = 4.69, *p* < 0.001), primary sensory, CC4 (ΔFA = 0.034, *t* = 3.15, *p* = 0.005), and lastly prefrontal regions, CC1 (ΔFA = 0.029, *t* = 2.73, *p* = 0.02). Figure 4 illustrates this interaction using model estimated marginal means.

**Figure 3.**
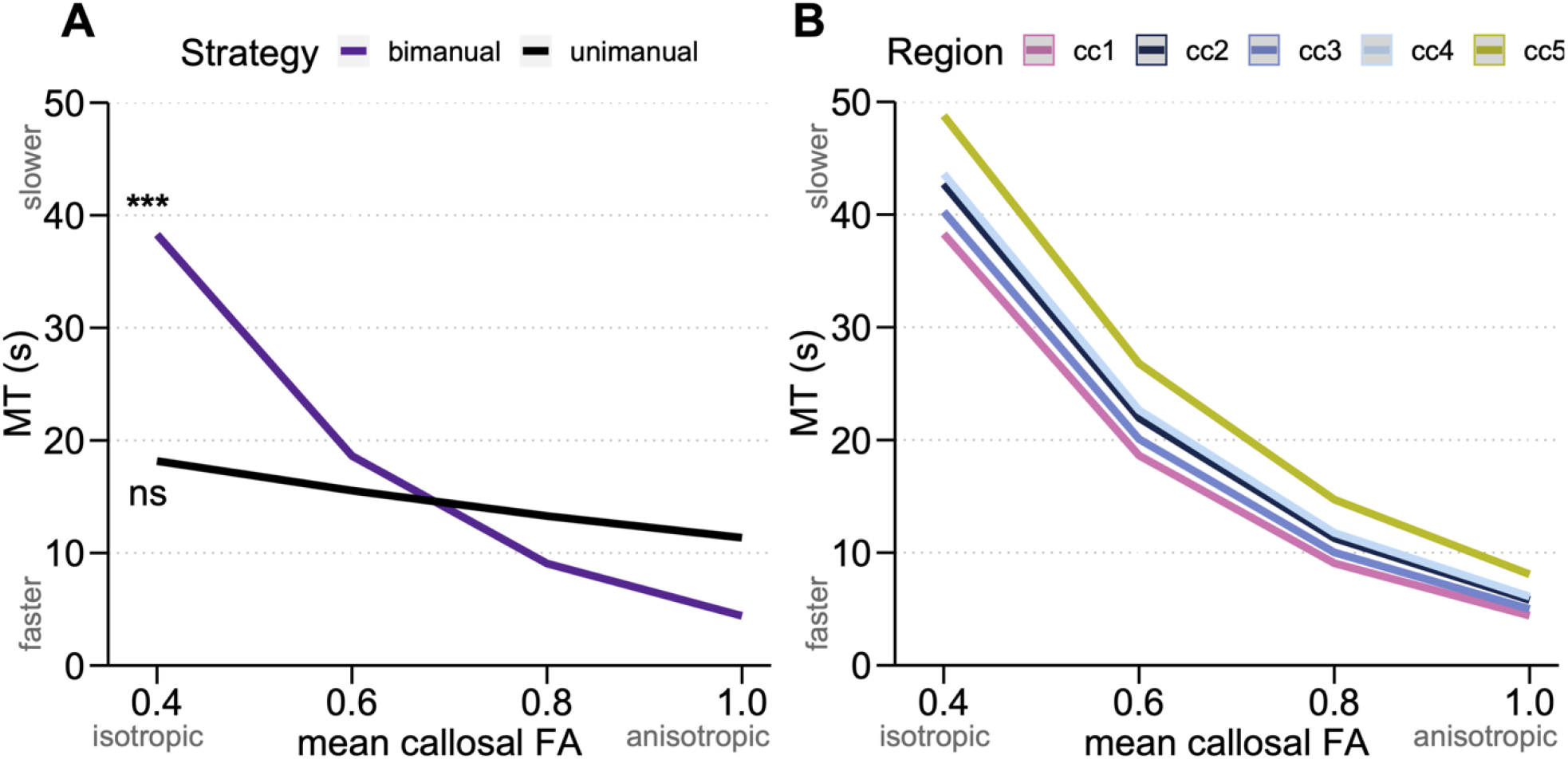
Movement time (MT) as a function of mean callosal FA as moderated by, A. strategy and B. callosal region. Lines are estimated marginal trends from the mixed effects model showing significant relationship between FA and MT for those who chose a bimanual but not unimanual strategy, and for the different CC regions.

**Figure 4.**
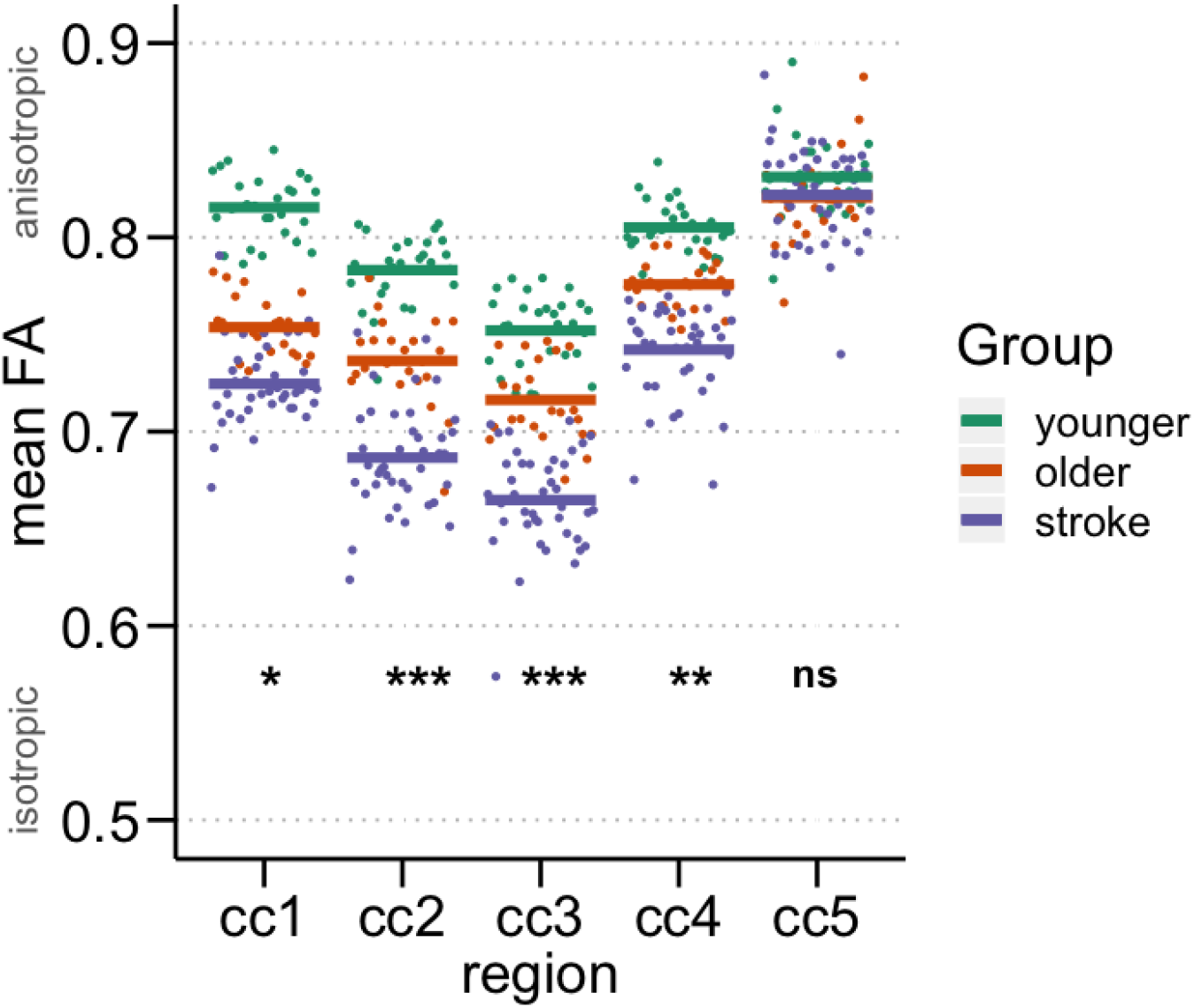
Model estimated marginal means for CC FA across the five regions along with individual data points. * p < 0.05, ** p < 0.01, *** p < 0.001.

## DISCUSSION

Our primary finding suggests that poor microstructural status of not only the sensorimotor fibers of the callosal body but also the non-sensorimotor regions of the anterior body and genu is associated with slower performance on a bimanual task in chronic stroke survivors. This relationship was found to be significant only for those who chose a bimanual strategy and not in those who chose a unimanual strategy, lending support to the idea that callosal fiber organization is uniquely important for interhemispheric communication underlying *bimanual* performance and is not simply a reflection of unimanual weakness and disuse.

A novel demonstration in this study is the relationship between CC2, the rostral portion of the body of the CC that connects homotopic premotor and supplementary motor regions, and bimanual performance in stroke. The medial wall of the frontal lobe has been shown to be causally involved in the control of self-initiated, self-paced cooperative bimanual movements ^21–24^. Patients who had undergone anterior callosotomy were significantly slower and the timing of movement initiations with both limbs were more imprecise ^25^ than in controls.

Preliowsky inferred that this may be because the anterior CC serves as a more direct route for the sharing of motor corollary discharges in the frontal lobe, enabling faster bimanual performance. The bimanual task we examined was cooperative, and participants were observed covertly as they self-initiated a movement strategy and completed the task at self-selected speed. Although not as extreme as callosotomy, it stands to reason that poor microstructural status of the anterior callosum in stroke survivors, delays transcallosal exchange of corollary discharges and slows performance on such a task.

Our secondary analysis revealed that compared to neurologically intact controls, chronic stroke survivors showed significant reduction in FA across the genu (CC1) and body (CC 2, 3, 4) of the corpus callosum, but no difference was found in the splenium (CC5). Given that the CC was not directly lesioned by the stroke, this widespread reduction in FA across the CC is evidence for transcallosal diaschisis. We extended previous findings of lower callosal FA in acute stroke (Pinter et al., 2020) survivors to the chronic phase, in those with mild to moderate motor impairment (Hayward et al., 2017; Stewart, Dewanjee, et al., 2017). Reductions in FA in the primary motor (CC3) and primary somatosensory (CC4) regions of the CC were less surprising and generally consistent with previous reports, with fairly similar effect sizes (ΔFA ∼0.05) to those reported in mild-to-moderate chronic stroke survivors.^5^ Changes observed in CC1 and CC2 were especially interesting as they suggest that transcallosal reorganization after stroke not only impacts the primary motor region, but also constituent regions of the larger sensorimotor networks, involved in the control of complex motor actions that require anticipatory motor planning and sequencing.

This study has several limitations. First, control imaging datasets were acquired on a different scanner; however, acquisition parameters were comparable, and scanner-related variability was considered as a random effect in the mixed-effect model. We were also able to validate our measure of FA by reproducing values similar to those reported previously.

Second, isotropy in FA values can result from a phenomenon known as fiber crossing; however, this issue is less pronounced for relatively medium distance fibers like that of the CC. Third, a very small subset of bimanual tasks was selected from a known large repertoire of bimanual skills, thus findings need to be replicated for other tasks requiring interlimb coordination. The assumption that the task studied here is a reliable proxy of interlimb coordination as studied in laboratory settings is a weak one, especially due to the lack of behavioral data in neurologically intact controls. While more ecologically valid and generalizable to bimanual performance in the real world, we are unable to definitively conclude details regarding the nature of interlimb coordination involved in such task. Finally, retrospective design and a relatively small sample size, especially of those individuals who chose a unimanual strategy, are also concerns. Whereas correlational analysis is the current standard in research using structural imaging for brain-behavior analysis, future work that extends structural diffusion imaging to multi-modal imaging including functional MRI along with larger samples and a prospective design might reveal new insights into transcallosal diaschisis after stroke in humans.

## CONCLUSION

Findings of this study lead us to conclude that in mild-to-moderate chronic stroke survivors with relatively localized lesions to the motor areas, callosal microstructure can be expected to change not only in the primary sensorimotor region, but also in the premotor, supplementary motor and prefrontal regions. A novel finding in this study was that these remote widespread changes in the callosal genu and body are associated with slower performance on cooperative bimanual tasks that require precise and interdependent coordination of the hands. Measures of callosal microstructure may prove to be a useful predictor of real-world bimanual performance in chronic stroke survivors and should be explored further in future investigations.

## Acknowledgements

This research study is supported by the National Institutes of Health under award numbers: NICHD F31HD098796 to R.V., NICHD R01HD065438 to C.W. and N.S., NINDS R56NS100528 to N.S., and NINDS R21NS120274 to N.S.

Tae-Ho Lee (Virginia Tech) and Mara Mather (USC Gerontology) for sharing DTI acquisition parameters and participant age for their OpenNeuro dataset. Members of the Neuroplasticity and Neurorehabilitation laboratory (USC Chan Division of Occupational Science and Occupational Therapy) for helpful discussions.

## Data and code availability

Data table and code for analysis are available in the first author’s OSF repository: <u>osf.io/7j9xe</u>

## References

1. von Monakow C. Die Lokalisation Im Grosshirn Und Der Abbau Der Funktion Durch Kortikale Herde. Vol LXIII. JF Bergmann; 1914. doi:10.1001/jama.1914.02570090083033

2. Li Y, Wu P, Liang F, Huang W. The microstructural status of the corpus callosum is associated with the degree of motor function and neurological deficit in stroke patients. PLoS ONE. 2015;10(4):1–17. doi:10.1371/journal.pone.0122615

3. Wang LE, Tittgemeyer M, Imperati D, et al. Degeneration of corpus callosum and recovery of motor function after stroke: A multimodal magnetic resonance imaging study. Hum Brain Mapp. 2012;33(12):2941–2956. doi:10.1002/hbm.21417

4. Stewart JC, O’Donnell M, Handlery K, Winstein CJ. Skilled Reach Performance Correlates with Corpus Callosum Structural Integrity in Individuals with Mild Motor Impairment after Stroke: A Preliminary Investigation. Neurorehabil Neural Repair. 2017;31(7):657–665. doi:10.1177/1545968317712467

5. Stewart JC, Dewanjee P, Tran G, et al. Role of corpus callosum integrity in arm function differs based on motor severity after stroke. NeuroImage Clin. 2017;14:641–647. doi:10.1016/j.nicl.2017.02.023

6. Lindenberg R, Zhu LL, Rüber T, Schlaug G. Predicting functional motor potential in chronic stroke patients using diffusion tensor imaging. Hum Brain Mapp. 2012;33(5):1040–1051. doi:10.1002/hbm.21266

7. Pinter D, Gattringer T, Fandler-Höfler S, et al. Early Progressive Changes in White Matter Integrity Are Associated with Stroke Recovery. Transl Stroke Res. 2020;11(6):1264–1272. doi:10.1007/s12975-020-00797-x

8. Hayward KS, Neva JL, Mang CS, et al. Interhemispheric Pathways Are Important for Motor Outcome in Individuals with Chronic and Severe Upper Limb Impairment Post Stroke. Neural Plast. 2017;2017. doi:10.1155/2017/4281532

9. Bonzano L, Tacchino A, Roccatagliata L, Abbruzzese G, Mancardi GL, Bove M. Callosal contributions to simultaneous bimanual finger movements. J Neurosci. 2008;28(12):3227–3233. doi:10.1523/JNEUROSCI.4076-07.2008

10. Gooijers J, Caeyenberghs K, Sisti HM, et al. Diffusion tensor imaging metrics of the corpus callosum in relation to bimanual coordination: Effect of task complexity and sensory feedback. Hum Brain Mapp. 2013;34(1):241–252. doi:10.1002/hbm.21429

11. Sisti HM, Geurts M, Gooijers J, et al. Microstructural organization of corpus callosum projections to prefrontal cortex predicts bimanual motor learning. Learn Mem. 2012;19(8):351–357. doi:10.1101/lm.026534.112

12. Caeyenberghs K, Leemans A, Coxon J, et al. Bimanual coordination and corpus callosum microstructure in young adults with traumatic brain injury: A diffusion tensor imaging study. J Neurotrauma. 2011;28(6):897–913. doi:10.1089/neu.2010.1721

13. Fling BW, Walsh CM, Bangert AS, Reuter-Lorenz PA, Welsh RC, Seidler RD. Differential callosal contributions to bimanual control in young and older adults. J Cogn Neurosci. 2011;23(9):2171–2185. doi:10.1162/jocn.2010.21600

14. Fling BW, Seidler RD. Fundamental differences in callosal structure, neurophysiologic function, and bimanual control in young and older adults. Cereb Cortex. 2012;22(11):2643–2652. doi:10.1093/cercor/bhr349

15. Franz EA, Eliassen JC, Ivry RB, Gazzaniga MS. Dissociation of Spatial and Temporal Coupling in the Bimanual Movements of Callosotomy Patients. Psychol Sci. 1996;7(5):306–310. doi:10.1111/j.1467-9280.1996.tb00379.x

16. Rowe JB, Toni I, Josephs O, Frackowiak RSJ, Passingham RE. The prefrontal cortex: Response selection or maintenance within working memory? Science. 2000;288(5471):1656–1660. doi:10.1126/science.288.5471.1656

17. Baxter MG, Parker A, Lindner CCC, Izquierdo AD, Murray EA. Control of response selection by reinforcer value requires interaction of amygdala and orbital prefrontal cortex. J Neurosci. 2000;20(11):4311–4319. doi:10.1523/jneurosci.20-11-04311.2000

18. Halsband U, Ito N, Tanji J, Freund H-J. The role of premotor cortex and the supplementary motor area in the temporal control of movement in man. Brain. 1993;116(1):243–266.

19. Sadato N, Yonekura Y, Waki A, Yamada H, Ishii Y. Role of the supplementary motor area and the right premotor cortex in the coordination of bimanual finger movements. J Neurosci. 1997;17(24):9667–9674.

20. Kornysheva K, Diedrichsen J. Human premotor areas parse sequences into their spatial and temporal features. Elife. 2014;3:e03043.

21. Preilowski BFB. Possible contribution of the anterior forebrain commissures to bilateral motor coordination. Neuropsychologia. 1972;10(3):267–277. doi:10.1016/0028-3932(72)90018-8

22. Brinkman J, Kuypers HGJM. Cerebral control of contralateral and ipsilateral arm, hand and finger movements in the split-brain rhesus monkey. Brain. 1973;96(4):653–674. doi:10.1093/brain/96.4.653

23. Kazennikov O, Hyland B, Wicki U, Perrig S, Rouiller EM, Wiesendanger M. Effects of lesions in the mesial frontal cortex on bimanual co-ordination in monkeys. Neuroscience. 1998;85(3):703–716. doi:10.1016/S0306-4522(97)00693-3

24. Stephan KM, Binkofski F, Halsband U, et al. The role of ventral medial wall motor areas in bimanual co-ordination. A combined lesion and activation study. Brain. 1999;122(2):351–368. doi:10.1093/brain/122.2.351

25. Eliassen JC, Baynes K, Gazzaniga MS. Anterior and posterior callosal contributions to simultaneous bimanual movements of the hands and fingers. Brain. 2000;123(12):2501–2511. doi:10.1093/brain/123.12.2501

26. Hawe RL, Sukal-Moulton T, Dewald JPA. The effect of injury timing on white matter changes in the corpus callosum following unilateral brain injury. NeuroImage Clin. 2013;3:115–122. doi:10.1016/j.nicl.2013.08.002

27. Voineskos AN, Rajji TK, Lobaugh NJ, et al. Age-related decline in white matter tract integrity and cognitive performance: A DTI tractography and structural equation modeling study. Neurobiol Aging. 2012;33(1):21–34. doi:10.1016/j.neurobiolaging.2010.02.009

28. Basser PJ, Pierpaoli C. Microstructural and physiological features of tissues elucidated by quantitative-diffusion-tensor MRI. J Magn Reson. 2011;213(2):560–570. doi:10.1016/j.jmr.2011.09.022

29. Varghese R, Kutch JJ, Schweighofer N, Winstein CJ. The probability of choosing both hands depends on an interaction between motor capacity and limb-specific control in chronic stroke. Exp Brain Res. 2020;(238):2569-2579. doi:10.1007/s00221-020-05909-5

30. Cirillo C, Brihmat N, Castel-Lacanal E, et al. Post-stroke remodeling processes in animal models and humans. J Cereb Blood Flow Metab. 2020;40(1):3–22. doi:10.1177/0271678X19882788

31. Xu J, Branscheidt M, Schambra H, et al. Rethinking interhemispheric imbalance as a target for stroke neurorehabilitation. Ann Neurol. 2019;85(4):502–513. doi:10.1002/ana.25452

32. Lotze M, Markert J, Sauseng P, Hoppe J, Plewnia C, Gerloff C. The role of multiple contralesional motor areas for complex hand movements after internal capsular lesion. J Neurosci. 2006;26(22):6096–6102. doi:10.1523/JNEUROSCI.4564-05.2006

33. Hoyer EH, Celnik PA. Understanding and enhancing motor recovery after stroke using transcranial magnetic stimulation. Restor Neurol Neurosci. 2011;29(6):395–409.

